# Exploring natural genetic variation in photosynthesis-related traits of barley in the field

**DOI:** 10.1101/2023.12.04.569890

**Authors:** Yanrong Gao, Merle Stein, Lilian Oshana, Wenxia Zhao, Shizue Matsubara, Benjamin Stich

**Affiliations:** Institute of Quantitative Genetics and Genomics of Plants, Heinrich Heine University, Düsseldorf, Germany; Institute of Bio- and Geosciences/Plant Sciences, Forschungszentrum Jülich, Jülich, Germany; Cluster of Excellence on Plant Sciences (CEPLAS); Julius Kühn Institute (JKI) – Federal Research Centre for Cultivated Plants, Institute for Breeding Research on Agricultural Crops, Sanitz, Germany; Xinjiang Seed Industry Development Center of China, Urumqi, China

**Author notes:** Corresponding author: Benjamin Stich, Benjamin Stich. E-mail address: Yanrong Gao Merle Stein Lilian Oshana Wenxia Zhao Shizue Matsubara.

**Keywords:** Barley, chlorophyll fluorescence parameters, crop yields, development, heritability, natural genetic variation, photosynthesis

## Abstract

Optimizing photosynthesis is considered an important strategy for improving crop yields to ensure food security. To evaluate the potential of using photosynthesis-related parameters in crop breeding programs, we measured chlorophyll fluorescence along with growth-related and morphological traits of 23 barley inbreds across different developmental stages in field conditions. The photosynthesis-related parameters were highly variable, changing with light intensity and developmental progression of plants. Yet, the variations in photosystem II (PSII) quantum yield observed among the inbreds in the field largely reflected the variations in CO_2_ assimilation properties in controlled climate chamber conditions, confirming that the chlorophyll fluorescence-based technique can provide proxy parameters of photosynthesis to explore genetic variations under field conditions. Heritability (*H*^2^) of the photosynthesis-related parameters in the field ranged from 0.16 for the quantum yield of non-photochemical quenching to 0.78 for the fraction of open PSII center. Two parameters, the maximum PSII efficiency in light-adapted state ( *H*^2^ 0.58) and the total non-photochemical quenching ( *H*^2^ 0.53), showed significant positive and negative correlations, respectively, with yield-related traits (dry weight per plant and net straw weight) in the barley inbreds. These results indicate the possibility of improving crop yield through optimizing photosynthetic light use efficiency by conventional breeding programs.

## Introduction

To satisfy the increasing demands for agricultural products at constant crop production areas, crop yields need to be increased by the year 2050 by about 25%-70% (Hunter *et al*., 2017). The potential genetic yield under an optimal environment is the product of four main factors: incident solar radiation, light interception efficiency, conversion efficiency, and harvest index (Bonington, 1977). The green revolution led to considerable increases of light interception efficiency and harvest index by introducing dwarfing genes into cereal crops (Hedden, 2003). However, some studies suggest that these two parameters are close to their theoretical maximum in modern crop varieties (e.g., Zhu *et al*., 2010). Accordingly, crop yield potential may be limited by the remaining bottleneck, the efficiency of light energy conversion by photosynthesis (source limitation) (Long *et al*., 2006*a*; Alvarez Prado *et al*., 2013; Kromdijk and Long, 2016). Thus, enhancing this conversion efficiency has become a breakthrough goal to improve crop yields (Zhu *et al*., 2010).

Notably, selection of yields might have unintentionally improved the conversion efficiency, as indicated by a positive relationship between photosynthesis and crop yields (Kromdijk and Long, 2016; Theeuwen *et al*., 2022). Still, the conversion efficiency has not reached the theoretical maximum in C_3_ plants (Long *et al*., 2006*b*; Zhu *et al*., 2010; Prosekov and Ivanova, 2018) after decades of selection for crop yields. This suggests that the selection for yields is not sufficient to fully explore and make better use of natural genetic variation of photosynthesis. Direct phenotyping and selection for photosynthesis parameters are needed to identify variations in photosynthetic capacity and source limitation of crop yield (Theeuwen *et al*., 2022).

Several studies have successfully increased yields through optimizing photosynthesis by genetic engineering (reviewed by Simkin *et al*., 2019), such as manipulating the Calvin– Benson cycle in wheat (Driever *et al*., 2017), carbon transport in rice (Gong *et al*., 2015) and soybean (Hay *et al*., 2017), or photoprotection in tobacco (Kromdijk *et al*., 2016) and in soybean (De Souza *et al*., 2022). However, the use of genetically modified crops is restricted in some parts of the world (Turnbull *et al*., 2021) and suggested yield improvements by the genetic modifications await rigorous tests in practical agricultural production conditions (Khaipho-burch *et al*., 2023). Classical breeding can offer an alternative or an additional approach. Indeed, natural variation of photosynthesis within (Wullschleger, 1993) and across species (Flood *et al*., 2011; van Bezouw *et al*., 2019; Garcia *et al*., 2022) can be exploited by classical breeding.

Natural genetic diversity of photosynthesis has been studied in cereals under field conditions. Driever *et al*. (2014) reported significant variations in photosynthetic capacity, biomass and yield in 64 wheat genotypes. Acevedo-Siaca *et al*. (2021*a*) observed high heritabilities for carbon assimilation-related parameters in 30 accessions of rice. However, the relationships between photosynthesis and yields observed in these studies were not consistent. For example, Carmo-Silva *et al*. (2017) observed a positive correlation between carbon assimilation rate and grain yields in field-grown wheat in pre- and post-anthesis stage, while Driever *et al*. (2014) found no correlation between carbon assimilation-related parameters and grain yield in field-grown wheat in pre-anthesis stages. A possible explanation for such discrepancies may be the dependency of the photosynthetic traits on environmental conditions and/or developmental stages of the plants, although further research is needed to clarify this. Furthermore, in barley, one of the most important cereal crops as well as a model for other cereals because of its simpler genetics, natural variation of photosynthesis has not been investigated under field conditions.

High-throughput phenotyping techniques are essential for investigating the natural genetic variation in photosynthesis. Photosynthesis is divided into two main processes, light reaction and CO_2_ assimilation, which can be assessed by chlorophyll fluorescence- and gas exchange-based techniques, respectively (Long *et al*., 1996; Baker, 2008). The analysis of chlorophyll fluorescence provides information on photosystem II (PSII) activity, such as the effective and the maximum quantum yields of PSII (Phi2 and Fv’/Fm’, respectively, for light-adapted state) or non-photochemical quenching (NPQ) (Baker, 2008). Measurement of gas exchange allows estimation of carbon assimilation rate (A) and related parameters (Sharkey, 2016). Recently, dynamic assimilation technique (DAT) was introduced to enable gas exchange measurements in non-steady state, which substantially increased the throughput compared to steady-state measurements (Saathoff and Welles, 2021) albeit still slower than chlorophyll fluorescence-based methods. For applications to crop breeding and selection, it is essential to check whether the genetic variations detected by these two techniques are comparable or not.

The objectives of this study were to 1) investigate genetic variation of photosynthesis-related parameters in barley across different developmental stages and evaluate the interaction between genotypes and environment in field conditions, 2) compare gas exchange- and chlorophyll fluorescence-based assessments of photosynthetic traits, and 3) assess correlation between photosynthesis-related and morphological or growth-related parameters. Based on the results obtained, we will consider the potential of using photosynthesis-related parameters in crop and particularly barley breeding programs.

## Materials and methods

### Field experimental design

Twenty-three spring barley (*Hordeum vulgare*) inbreds were selected from a world-wide collection of 224 barley landraces and cultivars based on their genetic and phenotypic diversity (Weisweiler *et al*., 2019). These 23 barley inbreds are the parents of the double round-robin population (Casale *et al*., 2022). All 23 inbreds were grown at three different locations (Bonn, Cologne, and Düsseldorf) in Germany in 2021. In Bonn and Düsseldorf, the experimental designs were alpha designs with two or three complete replications, respectively. In Cologne, two trials were performed, named in the following as mini-big plot and big plot. In the mini-big plot and big plot trials, the 23 inbreds were grown as replicated checks in an augmented design, where each inbred was replicated two times and one time, respectively. The plants were grown in 10 *m*^2^, 10 *m*^2^, and 2.25 *m*^2^ plots in Bonn, Cologne big plot, and Cologne mini-big plot, respectively. In Düsseldorf, single row plots with 33 kernels per row were used. Air temperature and precipitation were recorded during the field experiments in all three locations (Supplementary Fig. S1).

### Climate chamber experimental conditions and design

Based on the results of the field experiments, six representative barley inbreds (HOR1842, IG128216, IG31424, ItuNative, K10877, and W23829/803911) were selected for a climate chamber experiment. The experimental design was a randomized complete block design with three replicates. The growth conditions were as follows: 14 h/10 h light/dark photoperiod, 18°*C* /16°*C* temperature, and 55% relative humidity. The maximal light intensity measured at 15 cm from the light panel was 750 *μmol m*^−2^*s*^−1^.

### Assessment of photosynthesis-related parameters

In the field experiments, the top fully expanded leaves of three representative plants from each plot were measured from seedling stage (ZS13, (Zadoks *et al*., 1974)) to dough development (ZS87) using MultispeQ V2 device (Kuhlgert *et al*., 2016). We used the measurement protocol “Photosynthesis RIDES”, by which the intensity of actinic light was automatically set to the ambient light intensity measured by the built-in light sensor. The following parameters were used for the further analyses: liner electron flow (LEF), the fraction of open PSII centers (qL), the quantum yield of PSII (Phi2), the maximum efficiency of PSII in light-adapted state (Fv’/Fm’), the total NPQ (NPQt), the quantum yield of NPQ (PhiNPQ), the quantum yield of non-regulated dissipation processes (PhiNO), and relative chlorophyll content (SPAD). In addition, the MultispeQ also recorded environmental parameters, such as the intensity of photosynthetically active radiation (PAR), ambient temperature, ambient humidity, and ambient pressure.

In the climate chamber experiment, parameters of gas exchange were measured multiple times from tillering stage (ZS21) to dough development (ZS89) alongside the MultispeQ measurements. The measurements were made on the top fully expanded leaves on the main stem. Three different light intensities (PAR = 400, 800, 1500 *μmol m*^−2^*s*^−1^) were used as simulated low-light (LL), medium-light (ML) and high-light (HL) conditions for the MultispeQ measurements. Leaf-level gas exchange measurements were performed by LI-6800 (LI-COR Biosciences Inc., Lincoln). Three replicates per genotype were measured from 1 h after the onset of the light period. The settings inside the LI-6800 chamber were as follows: PAR was kept at 1500 *μmol m*^−2^*s*^−1^ (as in the simulated HL) with 50% blue and 50% red light, 400 *μmols*^−1^ air flow rate, 10,000 rpm fan speed, 55% relative humidity, and 18°*C* air temperature. The CO_2_ concentration inside the LI-6800 chamber was 400 ppm during pre-acclimation which lasted between 10 and 15 min. After the pre-acclimation, photosynthetic CO_2_ response (*A*/*C*_*i*_ ) curves were measured according to DAT (Saathoff and Welles, 2021). CO_2_ ramps were started from 1605 to 5 ppm with ramping rates of 200 ppm. The *A*/*C*_*i*_ curves were then analyzed using the “plantecophys” package (Duursma, 2015) in R version 4.0.3 to estimate the maximum rate of carboxylation ( *V*_*c,max*_ ), the maximum rate of electron transport ( *J*_*max*_ ), and triose phosphate utilization (*TPU*).

### Assessment of morphological and growth-related traits

To determine the relative growth rate (RGR) of the 23 barley inbreds, aboveground biomass data were collected in the field experiment in Düsseldorf at six different time points during the vegetation period: at 62, 69, 76, 83, 97, and 125 days after sowing (DAS). Plants of one row (initially 33 kernels were sown) per plot were harvested for the 23 genotypes with three replicate plots. Wild animals visited the trails and, thus, the number of damaged plants for each row was recorded.

The dry weight per row per plot was used to estimate the dry mass per plant (DMP), which was needed for the assessment of growth curve parameters, using the following equation:

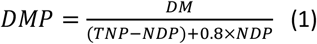

where TNP was the total number of plants, NDP the number of damaged plants, 0.8 was the completeness of the damaged plants based on the observation during the harvest. DMP calculated as described above, was corrected separately for each time point for replicate and block effects. The corrected values were then used for further analyses.

In the climate chamber experiment, the total aboveground DMP was directly measured by weighing at eight different time points (26, 36, 46, 57, 74, 102, 113, and 142 DAS) except for the two inbreds IG31424 and HOR1842, for which only the initial and the final DMP were determined at 26 and 142 DAS. Three replicates per genotype were collected for each time point.

To assess the relationship between DMP and time, logistic (Verhulst, 1838), power-low (Paine *et al*., 2012), and quadratic regression (Lithourgidis and Dordas, 2010) models were fitted. The quadratic regression model was used:

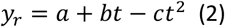

where *a* represents the initial biomass, *b* and *c* the growth rate parameters. This model had a high coefficient of determination (*R*^2^) and the highest heritability across all 23 barley inbreds. Thus, the quadratic regression was used for estimation of RGR. *RGR*_*a*_, *RGR*_*b*_, *RGR*_*c*_ represented the parameters in quadratic regression a, b, and c, respectively.

Morphological parameters were collected in multi-year and multi-environment field experiments that took place in the years 2017-2021 at Düsseldorf, Cologne, Mechernich, and Quedlinburg (Wu *et al*., 2022; Shrestha *et al*., 2022). Not all locations were used in all years to assess all parameters. Flag leaf length (FL, cm) and width (FW, cm), plant height (PH), flowering time (FT), awn length (AL, cm), spike length (EL, cm), and spikelet number in one row of the spike (SR), seed length (SL, mm), seed width (SW, mm), seed area (SA, mm^2^), and thousand grain weight (TGW, g), grain weight (GW, Kg/10 m^2^), and net straw weight (NSW, Kg/10 m^2^) were measured as morphological parameters. FL, FW, AL, EL were measured by ruler, SL, SW, and SA were measured by MARViN seed analyser (MARViNTECH GmbH, Germany), TGW was measured by MARViN and a balance.

The same set of morphological parameters was also measured in the climate chamber experiment. FL and FW were collected at 74 and 102 DAS with three replicates, and spike-related traits (AL, EL, SR, SL, SW SA, and TGW) were collected at 142 DAS with three replicates. Additionally, the total stem (without spike) weight per plant (SWP, g), total spike weight per plant (SKWP, g), total stem weight of main stem (TSWM, g), and spike weight of main stem (SKWM, g) were also collected in the climate chamber experiment. Harvest index (HI) was calculated using the following equation:

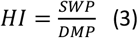

In addition, harvest index of main stem (MSHI) was calculated using the following equation:

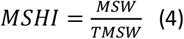

### Statistical analyses

#### Field experiment

Due to the strong dependence of photosynthesis on light intensity (Ogren, 1993), we considered three light intensity clusters when analyzing field measurements: LL, ML and HL conditions. These light intensity clusters were identified by K-means clustering of PAR and LEF. In addition, we also compared three main developmental phases of barley, i.e., slow expansion phase (SEP) (ZS<30), rapid expansion phase (REP) (30≤ZS<60), as well as anthesis and senescence phase (ASP) (ZS≥60). These two factors light intensity (L) and developmental phase (S), each with three levels, were considered when analysing the MultispeQ parameters from the field experiments based on the following linear model with the quantitative covariates light intensity (PAR) and developmental stage (ZS):

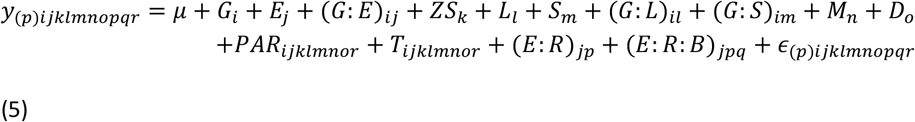

where *y*_(*p*)*ijklmnor*_ was the observed MultispeQ parameter across all light conditions and all developmental stages, *μ* the general mean, *G*_*i*_ the effect of the *i*^*th*^ inbred, *E*_*j*_ the effect of the *j*^*th*^ environment, (*G*: *E*)_*ij*_ the interaction between the *i*^*th*^ inbred and the *j*^*th*^ environment, *ZS*_*k*_ the effect of *k*^*th*^ zadok’s score of barley development, *L*_*l*_ the effect of *l*^*th*^ light intensity cluster, *S*_*m*_ the effect of *m*^*th*^ barely developmental phase, (*G*: *L*)_*il*_ the interaction between *i*^*th*^ inbred and *l*^*th*^ light intensity cluster, (*G*: *S*)_*im*_ the interaction between *i*^*th*^ inbred and *m*^*th*^ barley developmental phase, *M*_*n*_ the effect of the *n*^*th*^ MultispeQ device, *D*_*o*_ the effect of measurement date, (*E*: *R*)_*jp*_ the effect of the *p*^*th*^ replicate nested within *j*^*th*^ environment, (*E*: *R*: *B*)_*jpq*_ the effect of the *q*^*th*^ block nested within the *p*^*th*^ replicate in *j*^*th*^ environment, *PAR*_*ijklmnopqr*_ the light intensity of each measurement, *T*_*ijklmnopqr*_ the ambient temperature of each measurement, and *ϵ*_(*p*)*ijklmnopqr*_ the random error.

To estimate adjusted entry means for MultispeQ parameters of all inbreds, *G*_*i*_, *E*_*j*_, (*G*: *E*)_*ij*_, *ZS*_*k*_, *L*_*l*_, *S*_*m*_, (*G*: *L*)_*il*_, and (*G*: *S*)_*im*_ were treated as fixed effects, and *M*_*n*_, *D*_*o*_, (*E*: *R*)_*jp*_, (*E*: *R*: *B*)_*jpq*_ as random effects, *PAR*_*ijklmnopqr*_ and *T*_*ijklmnopqr*_ were covariates. Furthermore, we calculated adjusted entry means for all inbreds for each light intensity cluster as well as each developmental phase.

In addition, to evaluate the effect of each fixed factor and covariate, analysis of variance (ANOVA) was conducted.

To assess the heritability of each photosynthesis-related parameter at each developmental stage, which was considerably shorter than the above-mentioned three developmental phases, data were separated into eight stages from Zadok’s principal growth stages. The adjusted entry means were calculated based on the following model:

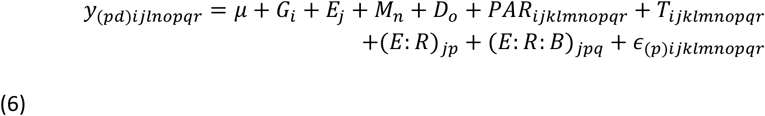

where, *y*_(*pd*)*ijlnopqr*_ was the photosynthesis-related parameter for each developmental stage across all other factors. Due to convergence problems, the interaction between *G*_*i*_ and *E*_*j*_ was removed from this model.

To assess the similarities among the barley genotypes with respect to their photosynthesis parameters, we performed hierarchical clustering by Ward’s minimum variance theory (Ward Jr, 1963) using the adjusted entry means of PSII parameters and SPAD at three different developmental phases. Furthermore, principal component analysis (PCA) was conducted by using the adjusted entry means calculated for each inbred in each of the developmental phase described before. The relationship between photosynthesis-related parameters and morphological or growth-related parameters of the inbreds was evaluated by Pearson correlation coefficients among adjusted entry means.

#### Climate chamber experiment

The adjusted entry means of carbon assimilation-related parameters from the climate chamber experiment were calculated based on the following model:

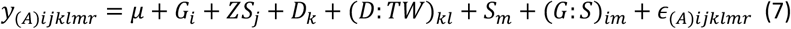

where *y*_(*A*)*ijklmr*_ was the carbon assimilation-related parameter, *D*: *TW*_*kl*_ the effect of the *l*^*th*^ time window in the *k*^*th*^ date of measurement, and *ϵ*_(*A*)*ijklmr*_ the random error. To estimate adjusted entry means for carbon assimilation-related parameters of six barley inbreds, *G*_*i*_, *ZS*_*j*_, *S*_*m*_ and (*G*: *S*)_*im*_ were treated as fixed effects, as well as *D*_*k*_ and *D*: *TW*_*kl*_ as random effects.

The relationship between photosynthesis-related parameters and morphological or growth-related parameters of the inbreds was evaluated by Pearson correlation coefficients between adjusted entry means.

#### Estimation of heritability

Broad-sense heritability ( *H*^2^ ) was estimated for both field and climate chamber experiments based on the following method:

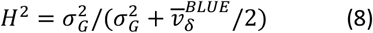

where 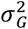 was the genotypic variance calculated based on the above models with a random effect for *G*_*i*_ and 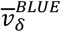 was the mean variance of the difference of two genotypic means (Holland *et al*., 2003; Piepho and Möhring, 2007).

To avoid the effect of the varying number of replicates, the *H*^2^ of photosynthesis-related parameters was estimated for each developmental stage based on the following equation:

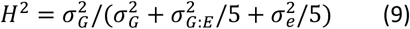

where 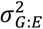 was the variance of the interaction of barley inbreds and environments, and 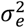 was the residual variance.

## Results

### Factors affecting photosynthesis-related parameters in the field

Parameters of PSII and SPAD were collected under field conditions with a wide range of PAR from 67 to 2172 *μmol m*^−2^*s*^−1^. In general, LEF increased as PAR increased, with an increasing variability among the individual observations at higher PAR (Fig. 1A). An increase of PAR was associated with a decrease of Phi2 and an increase of PhiNPQ, while PhiNO remained relatively stable (Supplementary Fig. S2A-C). Note that the sum of Phi2, PhiNPQ and PhiNO is equal to one. The light-dependent changes in Phi2 were accompanied by the corresponding changes in Fv’/Fm’ and qL (Supplementary Fig. S2D, E). The light response of NPQt was similar to that of PhiNPQ except that it often gave extreme values (Supplementary Fig. S2F). In contrast to these PSII parameters, SPAD values were not affected by momentary PAR (Supplementary Fig. S2G).

**Fig.1.**
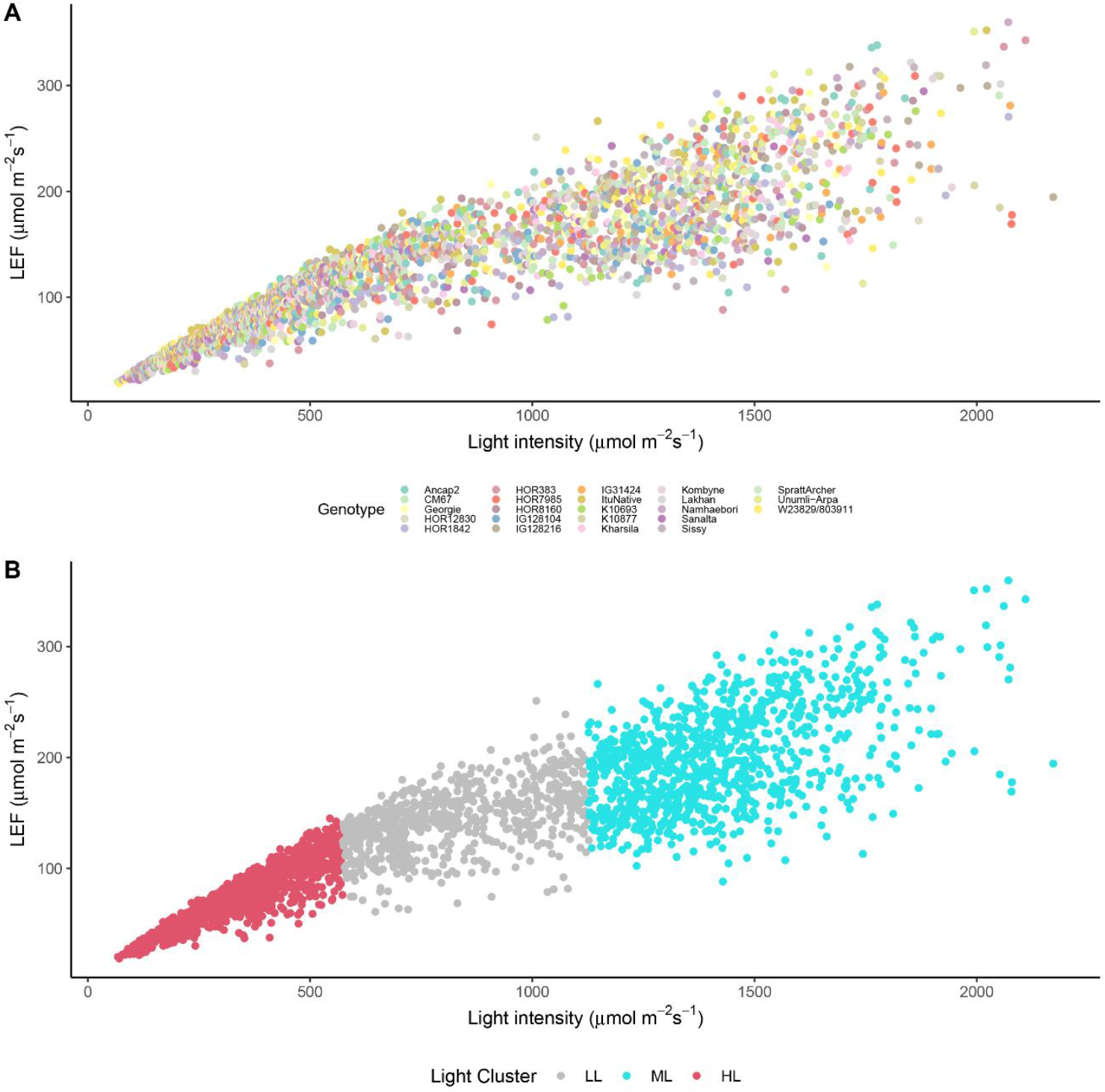
Liner electron flow (LEF) for 23 barley inbred lines in the field experiment in response to changing light intensity across all environments. The different colors of the dots in (A) indicate 23 different barley inbred lines. Three different colors of the dots in (B) represent the clusters of low (LL), medium (ML) and high (HL) light condition.

Given the strong influence of PAR on PSII parameters, K-means clustering was performed to separate the observations of LEF into three groups based on the light intensity: LL, ML, and HL conditions (Fig. 1B). As expected, significant (*P* < 0.05) differences in Phi2 and PhiNPQ, but not SPAD, were observed among the three light intensities (Fig. 2A). We then assessed the impact of developmental phase on these parameters: SEP, REP, and ASP (Fig. 2B). All three parameters showed significant (*P* < 0.05) differences between SEP and REP; Phi2 and SPAD increased from SEP to REP while PhiNPQ decreased (Fig. 2B). Thus, both PAR and developmental phases seem to affect Phi2 and PhiNPQ, whereas SPAD changed with development.

**Fig.2:**
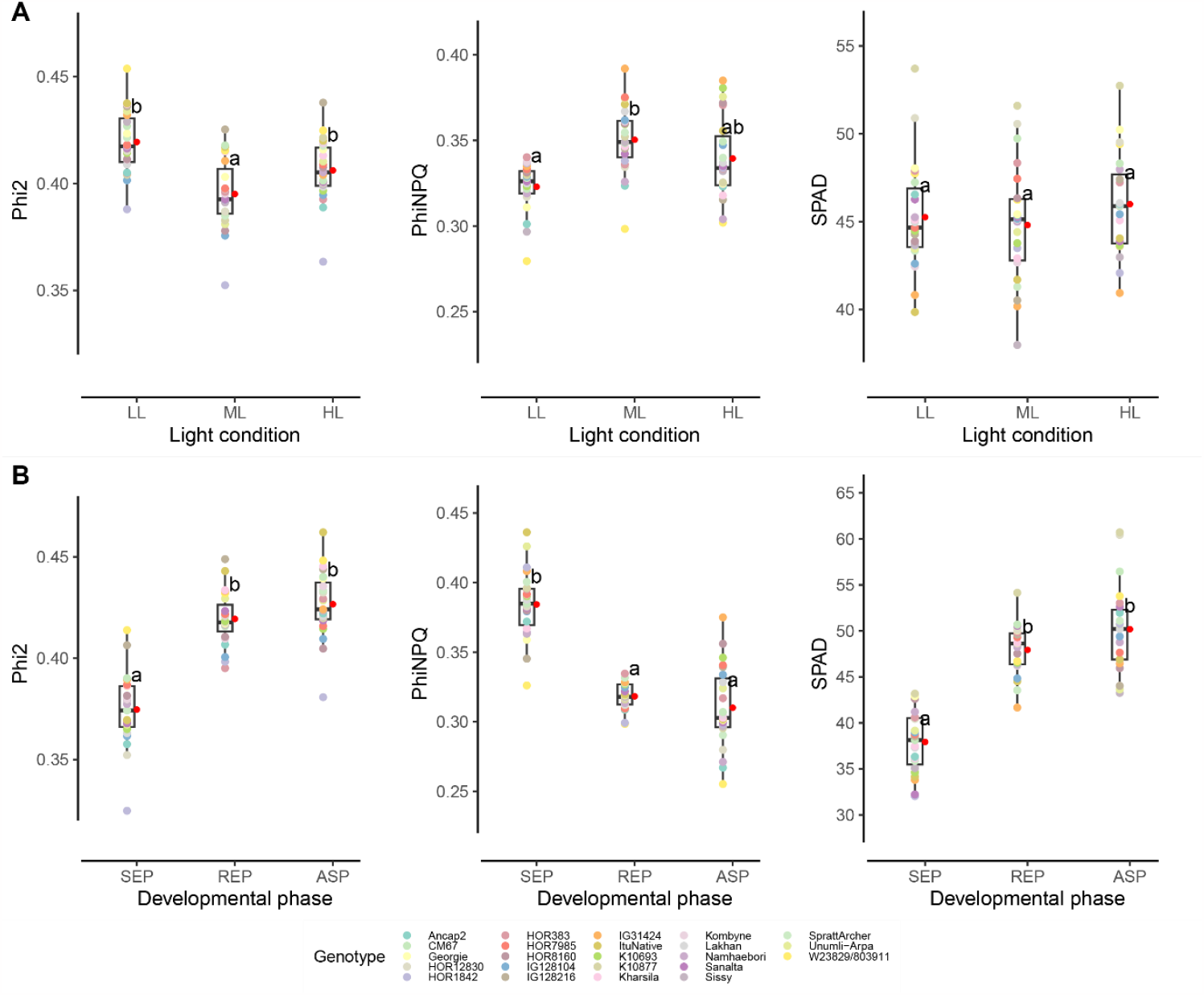
Effects of light intensity and developmental phase on the quantum yield of PSII (Phi2), the quantum yield of non-photochemical quenching (PhiNPQ) and relative chlorophyll content (SPAD) of 23 barley inbred lines. (A) Comparison of three light conditions: LL (low light), ML (medium light), and HL (high light). (B) Comparison of three developmental phases: SEP (slow expansive phase; Zadoks score, ZS, from 10 to 29), REP (rapid expansive phase; ZS from 30 to 59), and ASP (anthesis and senescence phase; ZS from 60 to 87). The colored dots represent the adjusted entry means for 23 barley inbred lines. The red point next to each box plot indicates the average across all inbreds for each light condition (A) or developmental phase (B). The letters next to each box plot indicate statistical significance. Different letters denote significant differences based on Tukey-test (P < 0.05) between the means for each parameter in each condition.

Significant ( *P* < 0.05 ) effects on PSII parameters and SPAD were observed for the interactions between genotype and environment 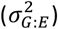, genotype and light condition 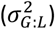 and genotype and developmental phase 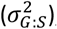, along with the effects of genetic variation 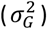) or other variables such as the date of measurement 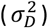, the used MultispeQ device 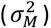, and the replicate 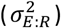 (Table 1). Analysis of variance also confirmed significant (*P* < 0.05) effects of PAR, light condition, developmental phase, and developmental stage (rated on the Zadoks scale) on the different PSII parameters and SPAD (Table 2). In addition, ambient temperature (T), which typically covaries with light intensity in the field, also significantly (*P* < 0.05) affected the PSII parameters except PhiNPQ in barley.

**Table 1:** Summary statistics of adjusted entry means, variance components, and broad-sense heritability (H^2^) for PSII parameters and SPAD measured in the field experiments. 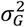 genotypic variance component; 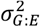 variance component of interaction between genotype and environment; 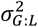 variance component of interaction between genotype and light conditions; 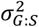 variance component of interaction between genotype and developmental phase; 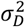variance component of date of measurement, 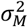variance component of MultispeQ device, 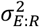 variance component of replicate in environment, 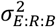 variance component of block nested within the replicate in environment. Note that the values of variance components of six photosynthesis-related parameters (Phi2, Fv’/Fm’, PhiNO, PhiNPQ, NPQt, qL) were multiplied with 10000.

| Trait | Mean | Min | Max | $\sigma_G^2$ | $\sigma_{G:E}^2$ | $\sigma_{G:L}^2$ | $\sigma_{G:S}^2$ | $\sigma_D^2$ | $\sigma_M^2$ | $\sigma_{E:R}^2$ | $\sigma_{E:R:B}^2$ | $\sigma_e^2$ | $H^2$ |
| --- | --- | --- | --- | --- | --- | --- | --- | --- | --- | --- | --- | --- | --- |
| LEF | 126.2 | 110.6 | 139 | 4.987 | 23.742 *** | 8.307 ** | 7.497 * | 151.189 *** | 236.731 *** | 0.325 | 2.935 | 469.69 | 0.42 |
| Phi2 | 0.41 | 0.37 | 0.43 | 0.90 *** | 0.84 *** | 0.17 | 0.24 | 16.25 *** | 12.54 *** | 0.72** | 0.1 | 21.29 | 0.74 |
| Fv'/Fm' | 0.67 | 0.64 | 0.7 | 0.5 | 0.54 ** | 0.71*** | 0.87*** | 11.73*** | 2.34*** | 0.52** | 0.03 | 25.52 | 0.58 |
| PhiNO | 0.26 | 0.22 | 0.3 | 1.36 *** | 0.62 ** | 0.65 ** | 0.48* | 6.99*** | 1.46 *** | 1.29*** | 0.06 | 34.12 | 0.74 |
| PhiNPQ | 0.34 | 0.29 | 0.37 | 0.09 | 1.37 *** | 0.84*** | 1.85*** | 32.62*** | 11.52*** | 0.29 | 0.17 | 32.57 | 0.16 |
| NPQt | 1.58 | 1.19 | 2.01 | 84.2 | 97.99*** | 168.23*** | 252.15*** | 1736.00*** | 372.24*** | 121.13*** | 7.58 | 5311.71 | 0.53 |
| qL | 0.35 | 0.28 | 0.42 | 4.25*** | 1.04* | 0.96* | 0 | 17.46*** | 5.51** | 3.38*** | 0.29 | 83.11 | 0.78 |
| SPAD | 45.35 | 40.65 | 52.68 | 2.091 | 5.850 *** | 0.544 | 5.309 *** | 33.570 *** | 31.471 *** | 1.199 * | 0.983 ** | 67.98 | 0.68 |

Asterisks indicate the significance of a likelihood ratio test (\*\*\*, \*\*, \* indicated $P < .001$ , .01, .05 respectively).

**Table 2:** Mean square values from analysis of variance for PSII parameters and SPAD measured in the field experiments. G is genotype, PAR is light intensity, T is ambient temperature, E is environment, L is light condition, ZS is Zadok’s score of barley development, S is development phases.

| Trait | G | PAR | T | E | L | ZS | S |
| --- | --- | --- | --- | --- | --- | --- | --- |
| LEF | 593.67 | 846205.72 *** | 14345.49 *** | 1751.45 * | 10979.05 *** | 951.84 | 1453.20 * |
| Phi2 | 0.48 * | 133.61 *** | 6.99 *** | 0.36 | 6.10 *** | 0.35 | 1.06 ** |
| Fv'/Fm' | 0.35 | 29.88 *** | 2.85 *** | 0.09 | 1.00 * | 4.74 *** | 2.71 *** |
| PhiNO | 0.73 * | 0.04 | 7.99 *** | 0.56 | 0.03 | 9.78 *** | 5.17 *** |
| PhiNPQ | 0.31 | 136.65 *** | 0.23 | 0.12 | 4.93 *** | 6.33 *** | 2.99 *** |
| NPQt | 69.53 | 5967.81 *** | 264.87 * | 10.22 | 86.6 | 686.60 *** | 394.44 *** |
| qL | 3.16 *** | 74.74 *** | 43.61 *** | 0.91 | 4.01 ** | 10.01 *** | 10.02 *** |
| SPAD | 94.54 | 0.19 | 216.02 | 45.77 | 80.4 | 4537.56 *** | 932.87 *** |

Asterisks indicate the significance of a likelihood ratio test (\*\*\*, \*\*, \* indicated $P < .001$ , .01, .05 respectively).

### Genetic variations in photosynthesis-related traits

When data from all light conditions and all developmental stages were combined together, *H*^2^ of the examined parameters ranged between 0.16 (PhiNPQ) and 0.78 (qL) (Table 1). Notably, when the heritability was calculated separately at different developmental stages as defined by Zadok’s growth scale, the *H*^2^ values of these parameters were considerably lower in the seedling growth stage and significantly (*P* < 0.05) higher in the dough developmental stage (Supplementary Fig. S3). In accordance, all PSII parameters and SPAD had low *H*^2^ values in the slow expansion phase (SEP) (Fig. 3). In the rapid expansion phase (REP), SPAD and Phi2 had the highest *H*^2^ while NPQt and Fv’/Fm’ had the lowest. In anthesis and senescence phase (ASP), the *H*^2^ of Phi2 decreased dramatically.

**Fig.3:**
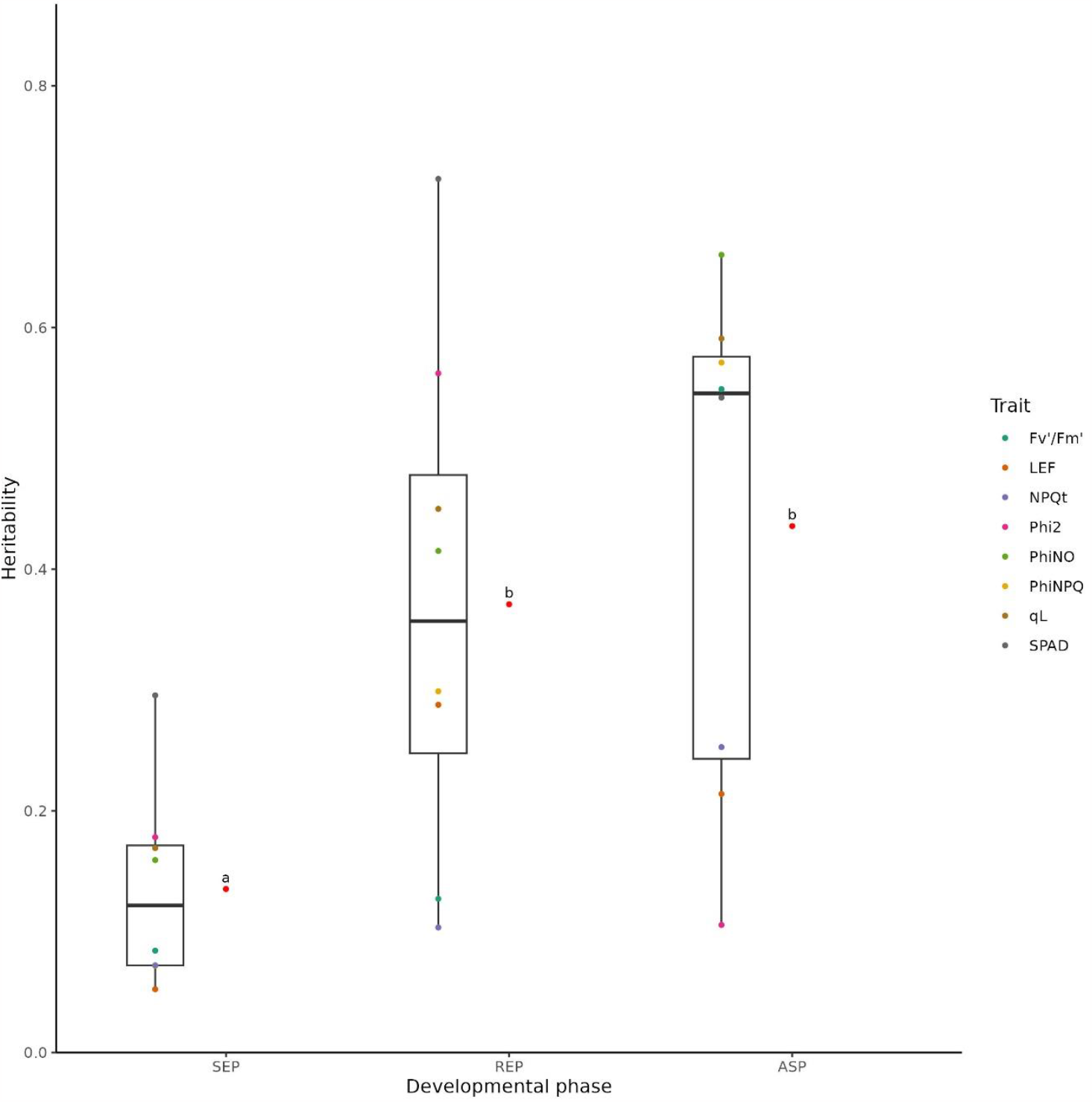
Heritability of PSII parameters and SPAD in different developmental phases. SEP: Slow expansive phase (Zadoks score (ZS) from 10 to 29); REP: Rapid expansive phase (ZS from 30 to 59); ASP: Anthesis and senescence phase (ZS from 60 to 87). The red point next to each box plot indicates the mean heritability across all traits for each developmental phase. Different letters next to each box plot indicate significant differences based on Tukey-test (P < 0.05) between the mean heritability.

Hierarchical cluster analysis of the adjusted entry means of the PSII parameters and SPAD observed in each of the three developmental phases indicated the presence of four major clusters among the 23 barley inbred lines (Fig. 4A). In general, the four clusters differed in the PSII parameters but not in SPAD (Fig. 5A). All PSII parameters except NPQt showed significant ( *P* < 0.05 ) differences among the four clusters in each of the three developmental phases (Fig. 5A).

**Fig.4:**
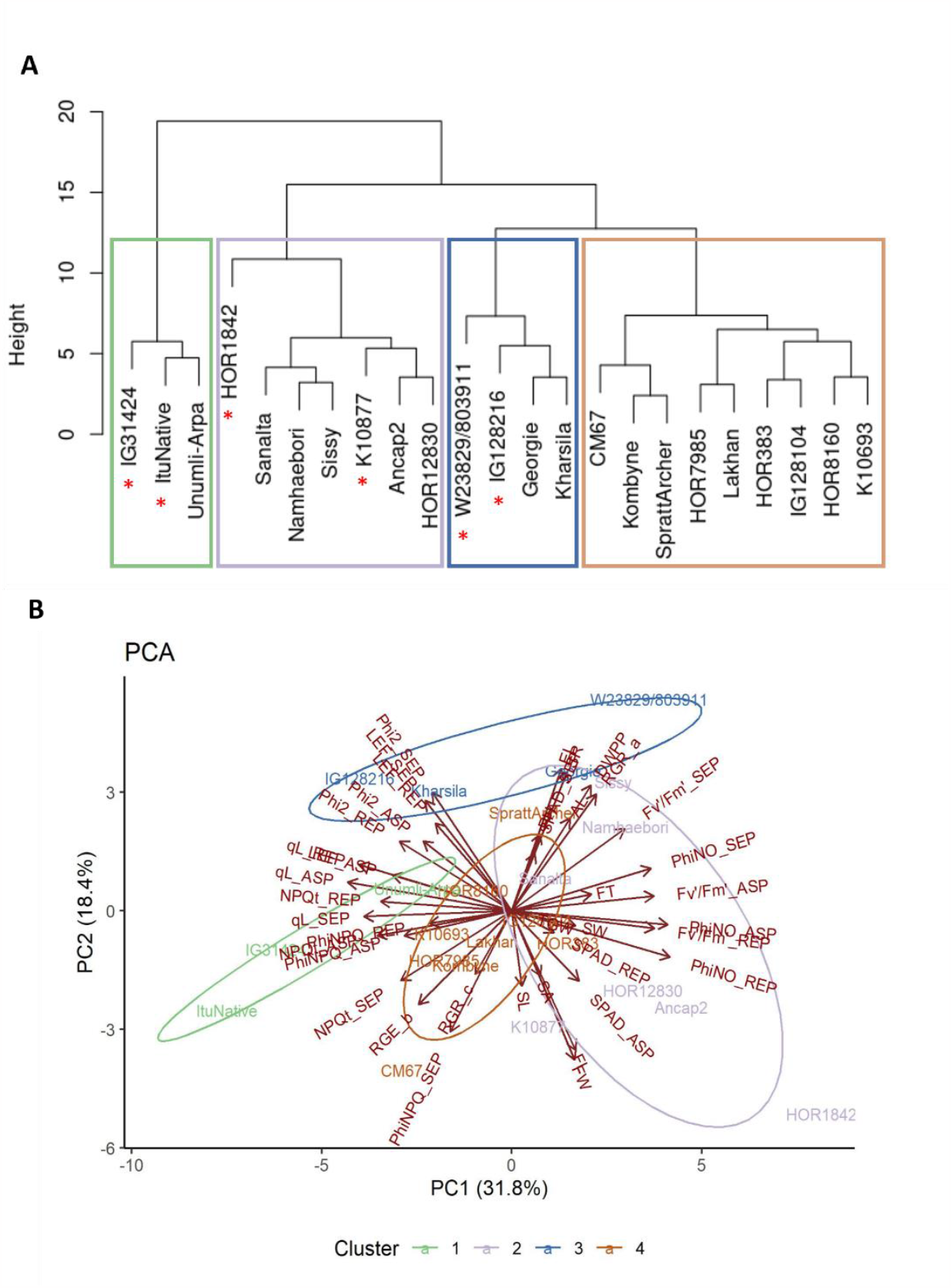
Hierarchical clustering (A) of 23 barley inbred lines based on their adjusted entry means for PSII parameters and SPAD in three developmental phases, and principal component analysis (B) based on the adjusted entry means of the combination of PSII parameters and SPAD in three developmental phases, the growth-related parameters based on dry mass per plant, and the morphological traits from multi-year and multi-environment experiments.

**Fig.5:**
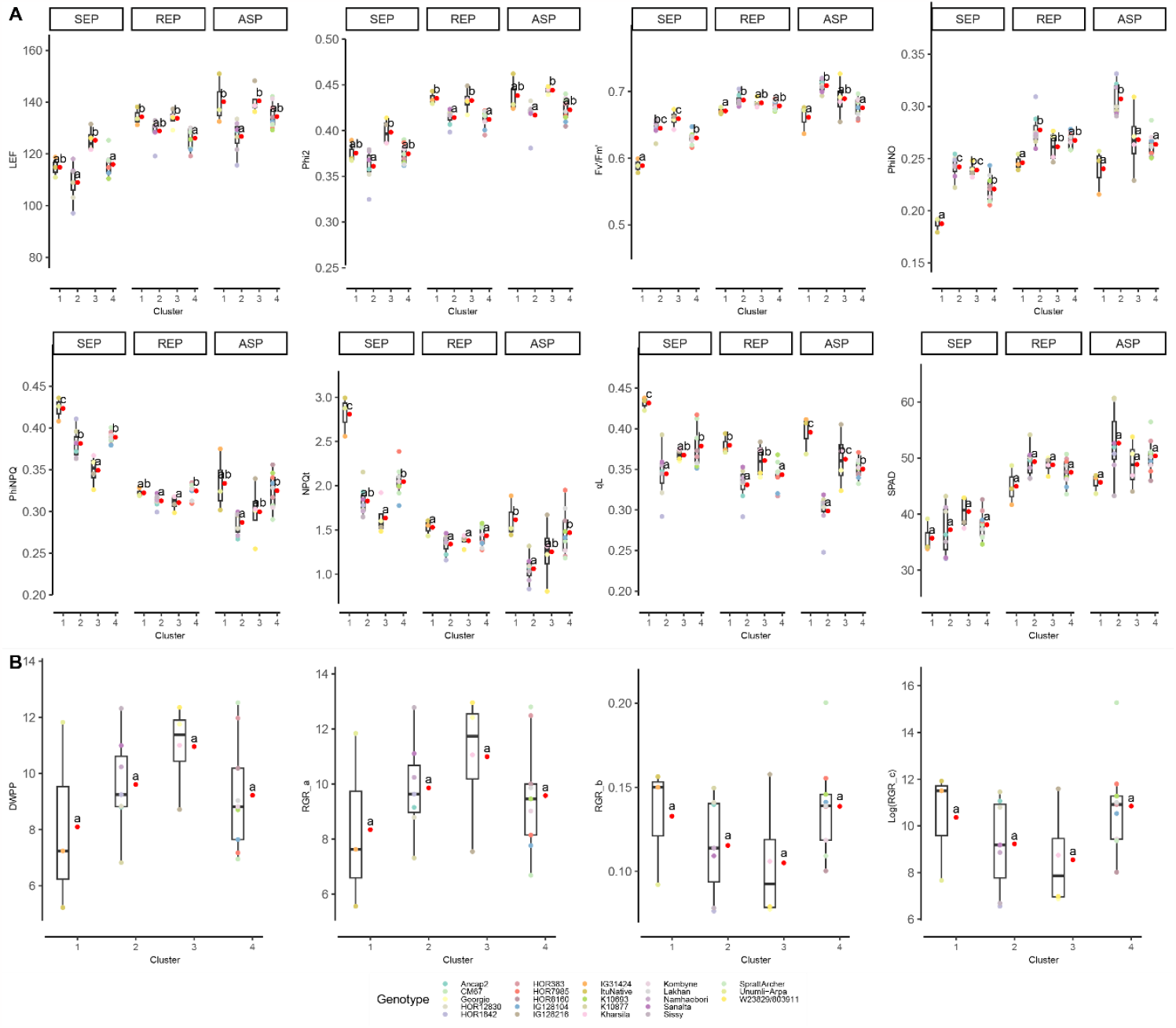
Comparison of PSII parameters, SPAD and growth-related parameters among the four clusters. (A) PSII parameters and SPAD in different developmental phases. (B) Dry mass per plant (DMP) and relative growth rates (RGR_a_, RGR_b_, RGR_c_) calculated from DMP based on the quadratic regression (y_r_ = a + bt − ct^2^). Due to the wide range of RGR_c_, log-transformed data of RGR_c_ was used. The red point next to each box plot indicates the mean of the parameters in each cluster. Different letters next to each box show significant differences based on Tukey-test (P < 0.05) between the clusters.

**Fig.6:**
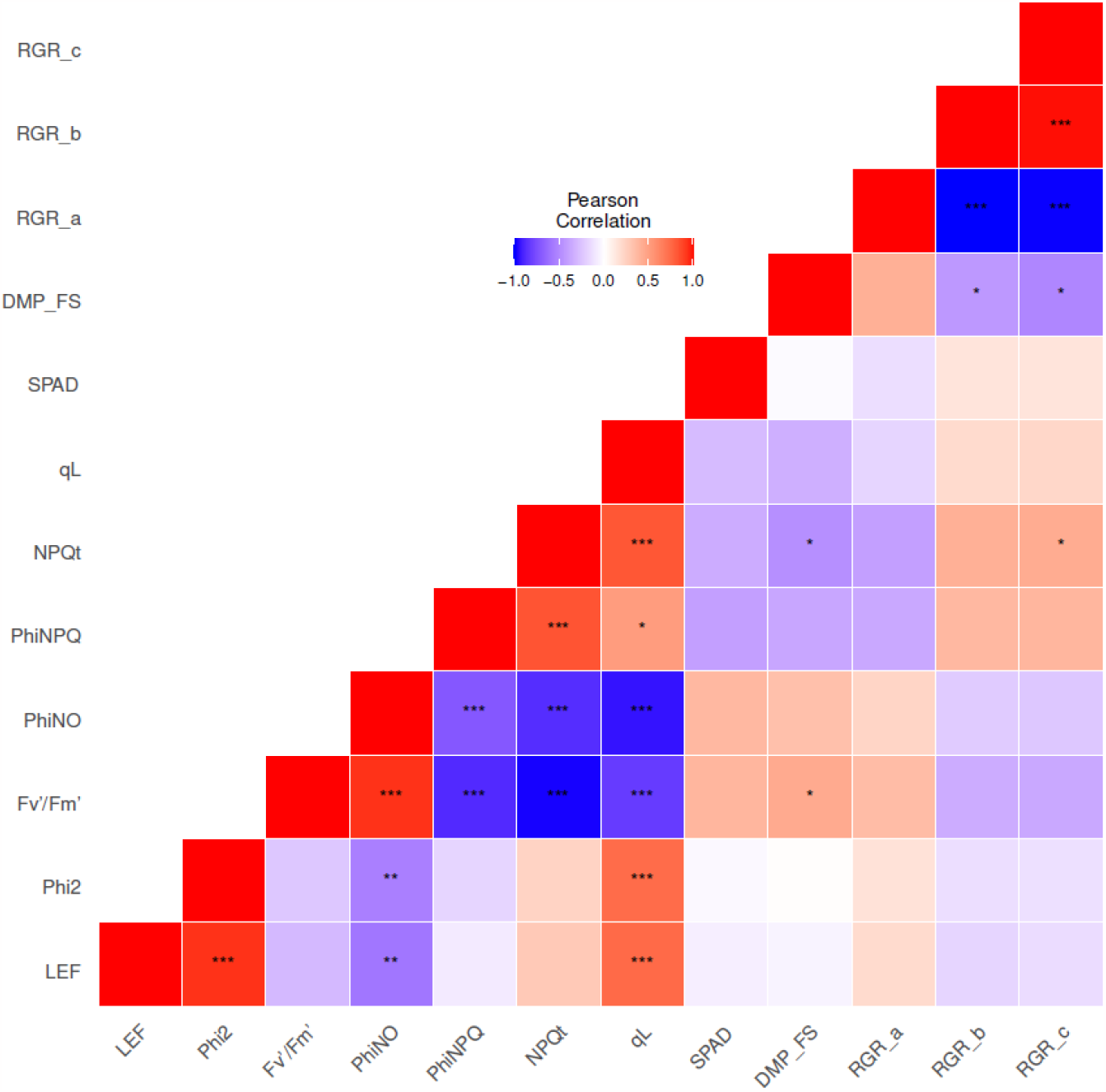
Person correlation coefficients calculated between pairs of adjusted entry means of 23 barley inbreds for photosynthesis- and growth-related parameters collected in the field. Asterisks indicate the significance level (***, **, * indicated P < .001, .01, .05 respectively).

We then asked whether the four clusters also represented differences in growth-related and morphological parameters among the 23 inbred lines. Relative growth rates (RGR) were calculated from the changes in DMP in the field experiment in Düsseldorf (Supplementary Fig. S4). Based on the growth parameters alone, the inbred lines could be divided into two clusters by hierarchical cluster analysis: DMP remaining at the same level after 100 days after sowing, and DMP increasing throughout the entire growth season (Supplementary Fig. S4). We observed no significant difference ( *P* = 0.3 ) in flowering time between the two groups. When PCA was performed on a combination of the PSII parameters and SPAD data shown in Fig. 2 as well as the growth-related parameters derived from Supplementary Fig. S4 and morphological traits collected from multi-year and multi-environment field experiments, the analysis revealed four clusters (Fig. 4B) that were very similar to those identified based on the PSII parameters and SPAD alone (Fig. 4A). Comparing the four clusters, we found no significant difference in DMP and RGR (Fig. 5B) or the morphological traits (Supplementary Fig. S5). Together, these results suggest that the clustering of the inbreds according to photosynthetic parameters primarily reflects genetic variations in photosynthetic traits among the 23 inbreds and is not confounded by differences in growth-related and morphological parameters.

### Comparison of gas exchange-based parameters and PSII parameters

Of the 23 barley inbreds, six (HOR1842, IG128216, IG31424, ItuNative, K10877, and W23829/803911) were selected to assess carbon assimilation-related parameters in climate chamber conditions. These six inbreds differed in the PSII parameters and SPAD in the field conditions. HOR1842 had the lowest adjusted entry means for Phi2, qL, and SPAD, IG128216 had the highest Phi2 and LEF. IG31424 had the highest PhiNPQ and the lowest SPAD. ItuNative had the lowest SPAD with relatively high Phi2, whereas K10877 had the highest SPAD with average values of PSII parameters. W23829/803911 was characterized by the lowest PhiNPQ and NPQt.

The gas exchange analysis in the climate chamber resulted in high *H*^2^ values for carbon assimilation-related parameters, ranging between 0.820 and 0.895 (Table 3). We observed a significant genetic variation (*P* < 0.05) for carbon assimilation at saturating light intensity (*A*_*sat*_) among the six inbreds (Fig. 7A); the adjusted entry means of *A*_*sat*_ were ranging from 14.7 (IG31424) to 19.7 (K10877) *μmol m*^−2^*s*^−1^. The differences in the maximal carboxylation (*V*_*c,max*_ ) and electron transport rates (*J*_*max*_ ) as well as triose phosphate utilization capacity (*TPU* ) were also significant (*P* < 0.05) among the six inbreds (Fig. 7A). Similarly, Phi2 (measured at LL, ML and HL) and SPAD showed significant ( *P* < 0.05 ) differences among the six inbreds (Fig. 7B). Carbon assimilation-related parameters underwent significant (*P* < 0.05) changes across the developmental phases, all peaking in REP together with Phi2 and SPAD (Fig. 7C, D). The *H*^2^ values for the PSII parameters and SPAD were also generally high, including 0.92 for Phi2 and 0.96 for SPAD (Table 3).

**Table 3:** Summary statistics of adjusted entry means, variance components and broad-sense heritability ( H^2^ ) for carbon assimilation-related parameters, SPAD and PSII parameters under 1500 μmol m^−2^s^−1^ assessed in the climate chamber experiment. 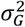 variance component of inbred; 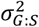 variance component of interaction between inbred and developmental phase; 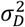 variance component of date of measurement, 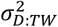 variance component of time window nested in date of measurement. Note, the values of variance components of six PSII parameters (Phi2_1500, Fv’/Fm’ _1500, PhiNO_1500, PhiNPQ_1500, NPQt_1500, qL_1500) were multiplied with 100.

| Trait | Mean | Min | Max | $\sigma_G^2$ | $\sigma_{G:S}^2$ | $\sigma_D^2$ | $\sigma_{D:TW}^2$ | $\sigma_e^2$ | $H^2$ |
| --- | --- | --- | --- | --- | --- | --- | --- | --- | --- |
| $V_{c,max}$ | 43.3 | 37.5 | 49.8 | 14.9 * | 6.432 * | 21.331 ** | 6.432 * | 50.886 | 0.848 |
| $J_{max}$ | 111.6 | 94.8 | 132.4 | 141.76 ** | 38.85 | 193.515 *** | 38.85 | 324.241 | 0.895 |
| $TPU$ | 7.61 | 6.37 | 9.13 | 0.717 ** | 0.23 * | 0.688 *** | 0.23 | 1.297 | 0.892 |
| $A_{sat}$ | 17.2 | 14.7 | 19.7 | 2.169 * | 1.32 * | 2.544 ** | 1.319 | 9.608 | 0.82 |
| LEF_1500 | 185.18 | 73.39 | 278.13 | 132.69 ** | 61.39 *** | 197.73 *** | 87.64 *** | 707.85 | 0.92 |
| Phi2_1500 | 0.27 | 0.11 | 0.41 | 0.03 ** | 0.01 *** | 0.04 *** | 0.02 *** | 0.16 | 0.92 |
| Fv/Fm_1500 | 0.6 | 0.21 | 0.75 | 0.05 ** | 0.02 ** | 0.05 ** | 0.02 ** | 0.26 | 0.92 |
| PhiNO_1500 | 0.23 | 0.04 | 0.45 | 0.03 ** | 0.01 | 0.05 *** | 0.02 * | 0.27 | 0.86 |
| PhiNPQ_1500 | 0.49 | 0.24 | 0.79 | 0.08 ** | 0.02 ** | 0.08 ** | 0.05 *** | 0.39 | 0.93 |
| NPQt_1500 | 2.33 | 17.89 | 0.64 | 11.70 * | 5.45 ** | 11.51 * | 6.32 ** | 80.99 | 0.9 |
| qL_1500 | 0.26 | 0.1 | 0.74 | 0.02 | 0.02 * | 0.11 *** | 0.02 * | 0.43 | 0.76 |
| SPAD | 44.19 | 6.28 | 75.47 | 36.22 *** | 3.81 * | 10.45 | 28.91 *** | 75.63 | 0.96 |

Asterisks indicate the significance of a likelihood ratio test (\*\*\*, \*\*, \* indicated $p_{value} \leq .001, .01, .05$ respectively).

**Fig.7:**
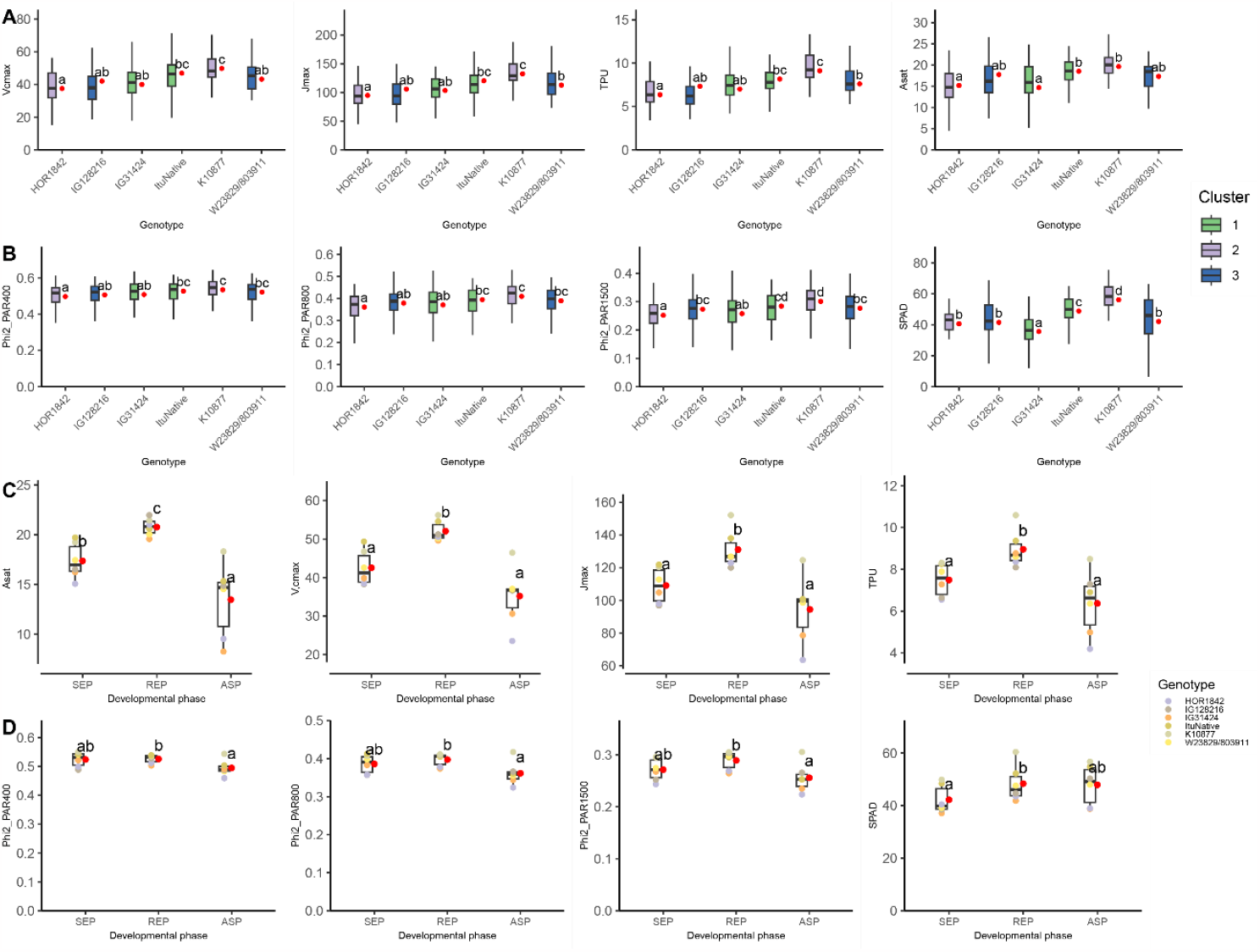
Comparison of the six barley inbred lines in the climate chamber. (A) Carbon assimilation-related parameters (A_sat_, V_c,max_, J_max_, TPU). (B) Phi2 under simulated LL (PAR 400 μmol m^-2^ s^-1^), ML (800 μmol m^-2^ s^-1^), and HL (1500 μmol m^-2^ s^-1^) conditions, and SPAD. For (A) and (B), the colors of boxes represent the clusters determined by the hierarchical clustering in Fig. 5a. (C) Adjusted entry means of A_sat_, V_c,max_, J_max_, TPU of the six inbred lines in SEP, REP, and ASP. (D) Adjusted entry means of Phi2 under the simulated LL, ML, and HL conditions, and SPAD of the six inbred lines in different developmental phases. The red dots next to boxes in (A) and (B) are the adjusted entry means of parameters of each genotype. The red dots next to boxes in (C) and (D) are the mean values of the six inbreds for each parameter in each developmental phase. Different letters next to each box denote significant difference based on Tukey-test (P < 0.05).

We observed significant (*P* < 0.05) positive correlations between SPAD, Phi2 and LEF (both measured at HL) and all four carbon assimilation-related parameters (determined at HL) in the climate chamber (Fig. 8). As anticipated, PhiNPQ and NPQt were negatively correlated with the four carbon assimilation-related parameters.

**Fig.8:**
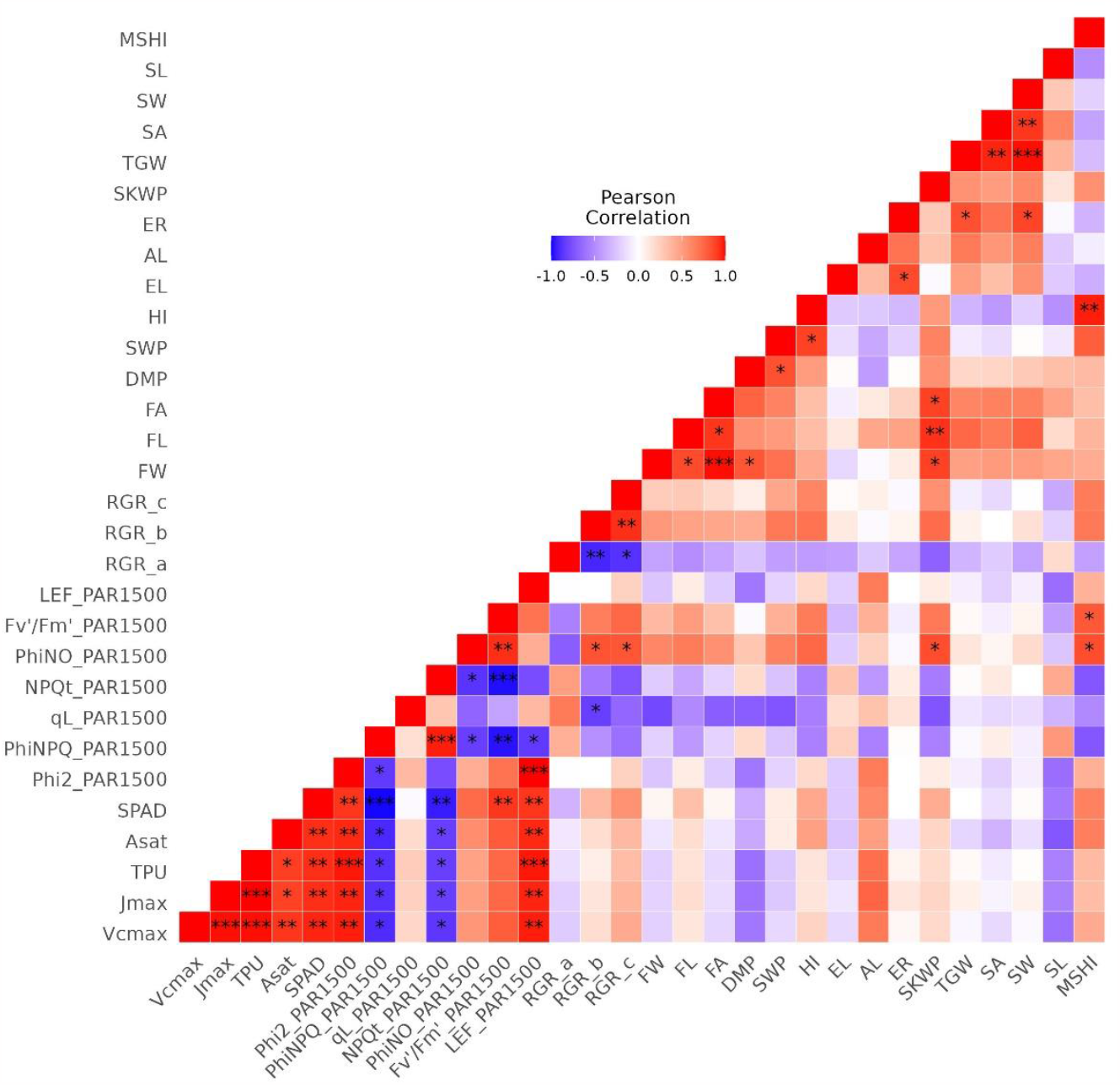
Person correlation coefficients calculated between pairs of adjusted entry means for six barley inbreds for photosynthesis-related parameters, dry mass per plant, harvest index (HI), flag leaf width (FW), flag leaf length (FL), flag leaf area (FA) and relative growth rate related parameters (RGR_a_, RGR_b_, RGR_c_), awn length (AL), spike length (EL), and spikelet number in one row of the spike (SR), seed length (SL), seed width (SW) seed area (SA) and thousand grain weight (TGW), total aboveground dry mass (DMP), total stem weight without spike weight (SWP), harvest index (HI), spike weight per plant (SKWP), main stem harvest index (MSHI) which were collected from the climate chamber experiments. Asterisks indicate the significance level (***, **, * indicated P < .001, .01, .05 respectively).

The adjusted entry means of the six barley inbreds showed significant (*P* < 0.05) positive correlations between Phi2 and carbon assimilation-related parameters in the climate chamber experiment (Fig. 9). In comparison, the correlations between these parameters assessed in the climate chamber experiment and Phi2 observed in the field were lower, with the highest correlation coefficient of 0.72 found for Phi2 at HL between these experiments.

**Fig.9:**
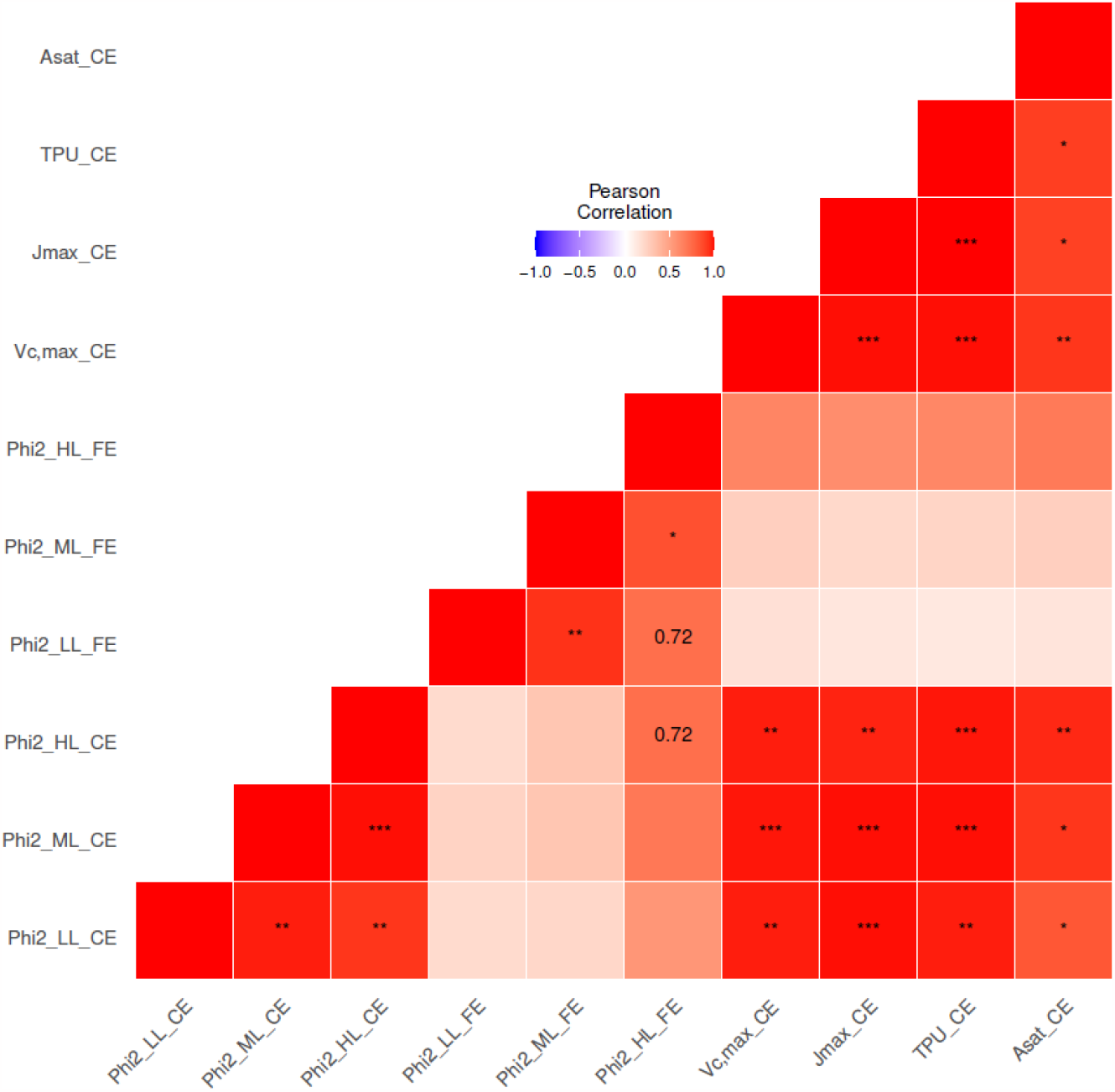
Person correlation coefficients calculated between pairs of adjusted entry means of six barley inbreds for Phi2 and carbon assimilation-related parameters measured in the climate chamber experiments (CE) and Phi2 measured in the field (FE). Phi2 was assessed separately for LL, ML and HL conditions. Caron assimilation was analysed at the light intensity of the HL condition. Asterisks indicate the significance level (***, **, * indicated P < .001, .01, .05 respectively).

### Relationship between photosynthesis-related parameters and growth or morphological parameters

Morphological parameters and DMP were determined in the climate chamber experiment to assess the relationship between the photosynthesis-related parameters and morphological or growth-related parameters. As done for the field experiments (Supplementary Fig. S4), RGR was calculated for the six inbred lines by fitting the DMP data to a quadratic regression (Supplementary Fig. S7). No significant correlation was observed between the morphological traits, DMP-based RGR and photosynthesis-related parameters among the six barley inbreds (Fig. 8).

We then made the same analysis using the data from the 23 inbred lines in the field experiments (Fig. 6). As expected, we found significant (*P* < 0.05) positive or negative correlations among the PSII parameters as well as among the growth-related parameters. No significant correlation was observed between SPAD and all PSII parameters when the adjusted entry means of all developmental stages and locations were considered (Fig. 6).

Comparing the PSII parameters and the growth-related parameters, the final DMP was significantly ( *P* < 0.05 ) positively and negatively correlated with Fv’/Fm’ and NPQt, respectively. RGR_c_ showed a significant (*P* < 0.05) positive correlation with NPQt (Fig. 6). Looking at the PSII parameters and morphological traits collected from multiple environments and years, significant ( *P* < 0.05 ) positive correlations were observed between two PSII parameters (PhiNO and Fv’/Fm’) and NSW (Fig. 10). In addition, significant negative correlations were observed between three PSII parameters (qL, NPQt, PhiNPQ) and NSW. Phi2, LEF and qL were significantly (*P* < 0.05) negatively correlated with flag leaf morphology (FL and FW) (Fig. 10).

**Fig.10:**
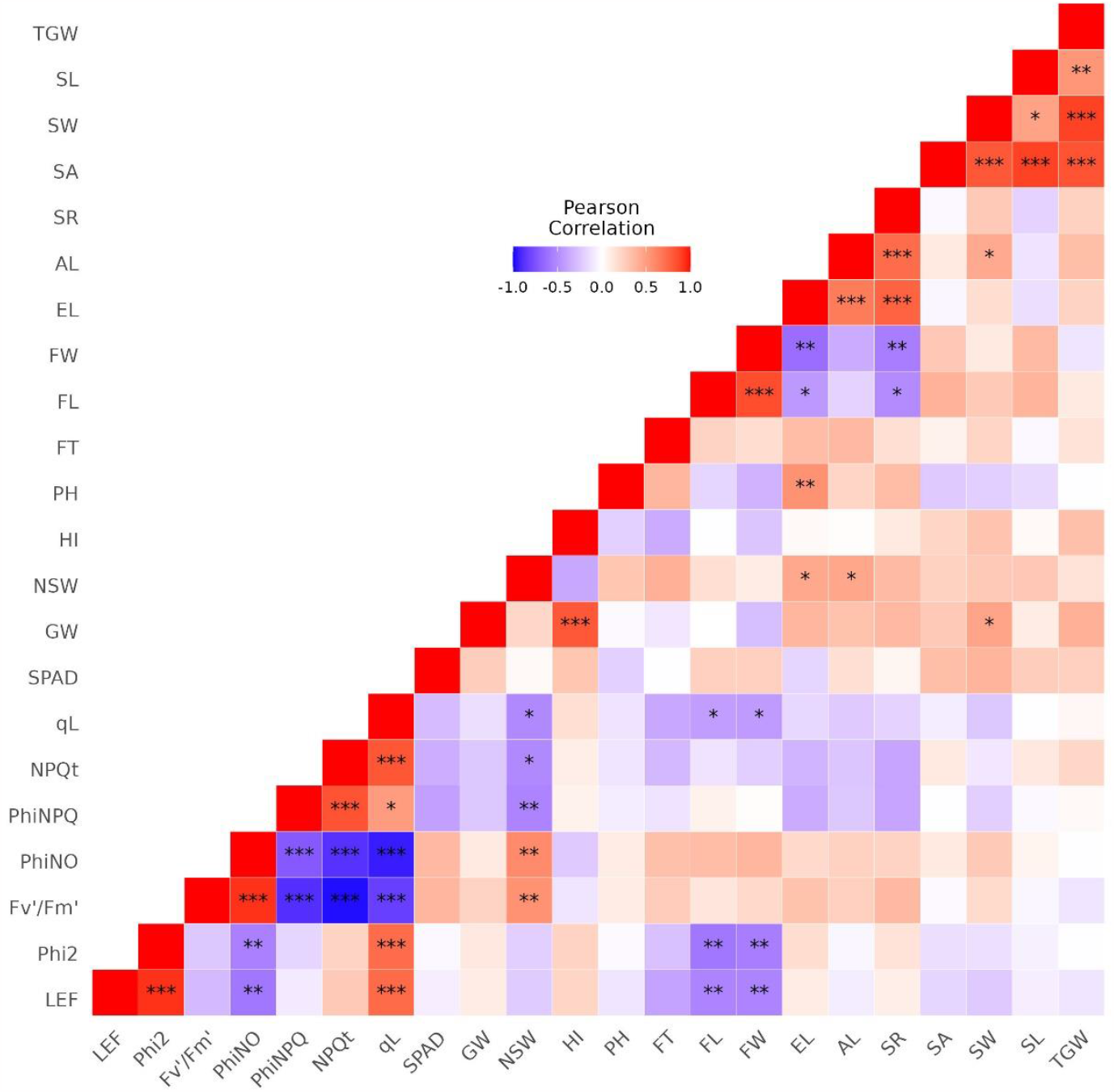
Person correlation coefficients calculated between pairs of adjusted entry means of 23 barley inbreds for PSII parameters, SPAD and morphological traits collected from multiple environments and years in the field conditions. Asterisks indicate the significance level (***, **, * indicated P < .001, .01, .05 respectively)

## Discussion

To meet the growing food demands, optimizing photosynthesis is a potential breeding target to support crop yield increases. In this study, we explored the natural genetic variation in photosynthesis-related parameters in 23 field-grown barley inbred lines.

### Comparison of chlorophyll fluorescence- and gas exchange-based techniques

In order to assess photosynthesis-related parameters in breeding programs, high-throughput methods are needed. This requirement is currently only fulfilled by chlorophyll fluorescence-based techniques. In comparison, detailed gas exchange measurements take, even with fast protocols of “DAT” (Saathoff and Welles, 2021), about 15 minutes for one measurement.

We evaluated the correlation between the genetic variations detected by chlorophyll fluorescence-based technique and by gas exchange-based technique. The climate chamber experiment showed high heritability for both photosynthesis-related parameters and carbon assimilation-related parameters (Tables 3), with significant (*P* < 0.05) genetic variance. There was a significant (*P* < 0.05) positive correlation between Phi2, which was assessed by chlorophyll fluorescence analysis, and carbon assimilation-related parameters in the climate chamber (Figs. 8 and 9).

The relationship between PSII electron transport and carbon assimilation has been investigated under laboratory conditions, such as in *Phaseolus vulgaris* L. (Farquhar *et al*., 1980), red campion, barley and maize (Genty *et al*., 1989) (for review see Bellasio et al., 2016). Our climate chamber experiment confirmed positive correlation between Phi2 and carbon assimilation-related parameters not only for one genotype but across diverse genotypes of barley. A similar correlation was observed across 41 spring wheat cultivars, in which the maximum quantum efficiency of PSII in dark-adapted state (Fv/Fm) was positively correlated with the assimilation rate measured in controlled environmental conditions (Sharma *et al*., 2015). The same group also showed that the genetic variation identified based on Fv/Fm (Sharma *et al*., 2012) was related to the difference in carbon assimilation rate (Sharma *et al*., 2015).

Despite the increasing number of studies focusing on photosynthesis under dynamic conditions (Keller *et al*., 2019; Acevedo-Siaca *et al*., 2021*c*; Fu and Walker, 2023; reveiwed by Long *et al*., 2022), few studies have demonstrated that the chlorophyll fluorescence-based parameters can be used to replace carbon assimilation parameters when investigating genetic diversity under natural environmental conditions. In this study, we observed a relatively high correlation coefficient (*r*=0.72) for Phi2 between the field and climate chamber experiments. The correlation between Phi2 in the field and carbon assimilation-related parameters in climate chamber experiments, however, was considerably lower (Fig. 9). Under field conditions, in which light intensity is changing dynamically, the balance between light reaction and carbon assimilation, as seen in a steady-state condition (Farquhar *et al*., 1980; Bellasio *et al*., 2016), may be broken (Rascher and Nedbal, 2006; Eberhard *et al*., 2008; Long *et al*., 2022). In particular, the lag of stomatal response to light intensity fluctuation limits CO_2_ uptake, resulting in a lower CO_2_ concentration inside the leaf. This will decrease carbon assimilation (Pearcy, 1990; Lawson *et al*., 2012) and increase photorespiration in species with C3 photosynthesis, eventually leading to reduced grain yields (Walker *et al*., 2016; Cavanagh *et al*., 2022).

Nevertheless, our observations in the present study, namely, 1) the significant positive correlation between Phi2 and carbon assimilation-related parameters in the climate chamber, 2) the positive correlation of Phi2 between the climate chamber and field conditions, and 3) the similar genotype ranks of Phi2 under both conditions, indicate that the chlorophyll fluorescence-based high-throughput technique can provide proxy parameters of photosynthesis to study genetic diversity under natural environmental conditions.

To use chlorophyll fluorescence-based techniques to assess highly variable photosynthesis parameters (Figs. 1 and 2) in breeding programs, however, it is important to consider the factors contributing to their variations.

### Factors contributing to photosynthesis variability in the field conditions

We observed high variability in photosynthesis-related parameters among 23 barley inbreds in the field (Table 1, Fig. 2). Four main factors are potentially contributing to the high variability of photosynthesis-related parameters: 1) environmental conditions, 2) developmental stages, 3) genetic diversity, and 4) interaction among genotypes, environment conditions and developmental stages. Below we will discuss these factors one by one.

#### 1) Growth environment of spring barley

We observed significant (*P* < 0.05) effects for the design variables, namely, location of the experiment, date of measurement (Table 1), as well as replicate 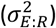. This can be explained by the dynamic environmental conditions during the growth season of spring barley, which was from late March to the beginning of August. Daily average temperature was fluctuating with an increasing trend during the growth season (Supplementary Fig. 1). Fluctuations in light intensity occurring within and between days must also have affected photosynthesis. We observed a significant (*P* < 0.05) effect of light intensity (PAR) and ambient temperature (T) (Table 2), which we considered as covariants because of the variability within a day and location of the measurement. In addition, lower temperature in May (Supplementary Fig. 1) might have suppressed Phi2 in SEP compared to the other two developmental phases (Fig. 2B) (Bagley *et al*., 2015; Moore *et al*., 2021).

In parallel to the changes of temperature and light intensity, photosynthesis efficiency typically exhibits diurnal (Flood *et al*., 2016) and seasonal patterns (Keller *et al*., 2019). Leaf movement (Flood *et al*., 2016), in interaction with dynamic environments, could also affect photosynthesis.

The significant (*P* < 0.05) variance components observed in our study for environmental factors, namely date (D), PAR and ambient temperature (T), indicate that single time point measurements are not sufficient to draw conclusions on genetic variation in photosynthesis under field conditions.

#### 2) Developmental stages

Most previous studies exploring the genetic diversity of photosynthesis in cereals focused on carbon assimilation in the flag leaf, which is the most important leaf in pre- or post-anthesis (Driever *et al*., 2014; Carmo-Silva *et al*., 2017; Acevedo-Siaca *et al*., 2021*b*). In our study, however, significant ( *P* < 0.05 ) differences in photosynthesis-related parameters were observed both in the field and climate chamber experiments at different developmental stages (Figs. 2 and 7C, D) (Tables 1 and2). As our analysis was corrected for environmental conditions, the observed differences in developmental phases are not due to the environmental changes during the experiments but due to the developmental stage of the plant itself. This is in accordance with the earlier reports of changing heritability for photosynthesis-related parameters across the lifespan of Arabidopsis (Flood *et al*., 2016) and changing QTLs (Meyer *et al*., 2021) detected for plant growth across the cultivation period under controlled conditions.

Photosynthesis-related parameters are not only linked to plant development, but also to leaf development (Wingler *et al*., 2004; Bielczynski *et al*., 2017). At the plant level, the sink tissues in SEP are mainly growing leaves and roots, while more new sink tissues emerged in REP, such as larger root system and formation of inflorescence meristem (Alqudah and Schnurbusch, 2017). The increased sink activity may explain the higher Phi2 in REP than SEP (Fig. 2b). In ASP, barley went from the anthesis stage to grain filling stage, which may further increase the sink strength. However, photosynthesis-related (source) parameters did not show corresponding increases in ASP compared to REP. In fact, it has been proposed that spike photosynthesis may serve as the major photosynthesis source for grain filling, as previously shown for wheat (Maydup *et al*., 2010; Vicente *et al*., 2018; Molero and Reynolds, 2020).

At the leaf level, photosynthesis efficiency is low in young growing leaves and increases gradually to reach the maximum during leaf expansion (Bielczynski *et al*., 2017). Subsequently, declining activity of photosynthesis after the anthesis period has been reported in many studies (Wingler *et al*., 2004; Liu *et al*., 2017; Miao *et al*., 2018; Yang *et al*., 2018). In the present study, we always took the measurements from the top fully expanded leaves throughout all developmental stages of the plants to minimize the effect of leaf development. As a result, no significant difference in Phi2 and SPAD was observed between REP and ASP in our experiments.

#### 3) Genotypic effect

In order to evaluate the potential of classical breeding to optimize photosynthesis, the relative importance of genotypic effects versus non-genotypic effects on photosynthesis-related parameters (i.e., the heritability) needs to be considered. The heritability varied between 0.16 and 0.78 (Table 1). The relatively high heritability together with the significant (*P* < 0.05) genetic variances found for Phi2, PhiNO, and qL suggest that these parameters could be suitable targets of photosynthesis breeding programs.

Notably, similar to the large variations in photosynthetic performance found across the whole growth season of barley, the heritability of photosynthesis-related parameters also varied in the different developmental stages of plants, with the lowest values in the seedling stage (Supplementary Fig. S3). This dynamic heritability suggests that the relative contributions of environment and genetics are not stable during the plant growth and development (Yang *et al*., 2015). Dynamic heritability was also observed in Arabidopsis (Flood *et al*., 2016) under controlled conditions. Unlike in our study, however, the dynamic heritability of that study was mainly attributed to the variation of genetics and diurnal changes of photosynthesis.

Taking the photosynthetic measurements when the heritability is low may not be efficient to select the optimal genotype (Visscher *et al*., 2008). Better knowledge about the dynamic pattern of heritability change during plant growth and development would help us select the developmental stage(s) when the genotype contributes most to the variation of photosynthesis, thus facilitating more targeted and efficient plant breeding (Flood *et al*., 2016).

#### 4) Interactions between genotype, environment and developmental stage

As discussed above, photosynthesis is highly responsive to environmental conditions and subject to developmental influences. We found significant (*P* < 0.05) interaction effects between genotype and environment (*G*: *E*), genotype and light condition (*G*: *L*), as well as genotype and developmental phases (*G*: *S*) on photosynthesis-related parameters (Table 1). Importantly, the interaction of *G*: *S* strongly suggests that it is necessary to carefully choose photosynthesis-related parameters in different developmental stages of the plants to explore genetic variation in crop breeding programs.

### Covariation between photosynthesis variability and yields related-traits

Having confirmed genetic variation for photosynthesis-related parameters in 23 spring barley inbreds, we also analyzed the relationship between photosynthesis- and yield-related traits.

The yield-related traits were collected in multi-environment and multi-year experiments, while photosynthesis-related parameters were collected in three different locations in one year. The genotype*environment interactions, which are important for not only yield but also photosynthesis, are most likely responsible for the non-significant correlations between photosynthesis-related parameters and yield-related traits in these experiments. In addition, it is difficult to connect leaf-level photosynthesis to plant-level biomass (Acevedo-Siaca *et al*., 2021*a*). Nevertheless, we observed significant (*P* < 0.05) positive correlations between Fv’/Fm’ and NSW and DMP_FS,_ as well as significant ( *P* < 0.05 ) negative correlations between NPQt and NSW and DMP_FS_, based on the adjusted entry means across all developmental stages (Supplementary Fig. S6). Thus, Fv’/Fm’ and NPQt might have the potential to be targeted in breeding programs for crop yield improvement.

## Conclusions

The results of this study show that chlorophyll fluorescence-based technique is able to detect genetic variation in PSII-related parameters in barley under climate chamber conditions, and Phi2, PhiNO and qL could be suitable parameters to detect genetic variation under field conditions. The significant effects of environmental factors and significant interactions between the genotype and environments indicate that the non-genetic effects must be taken into account when designing field studies. Significant correlations observed between photosynthesis-related traits (Fv’/Fm’ and NPQt) and yield-related traits (NSW, DMP_FS_) in barley under field conditions suggest the possibility of improving crop yields via optimizing photosynthesis through conventional breeding approach. If the breeding program is directly targeting at high photosynthesis efficiency, the rapid expansion phase is the developmental stage of choice to take measurements in barley because of the observed high heritability of photosynthesis-related parameters.

## List of photosynthesis-related parameters assessed in this study

Parameterh: Descriptionh
Fv’/Fm’: Maximum efficiency of PSII in light-adapted state
*J*_*max*_: Maximum rate of electron transport
LEF: Liner electron flow
NPQt: Total non-photochemical quenching
Phi2: Quantum yield of PSII
PhiNO: Quantum yield of non-regulated dissipation processes
PhiNPQ: Quantum yield of non-photochemical quenching
PSII: Photosystem II
qL: Fraction of PSII open center
SPAD: Relative chlorophyll content
TPU: Triose phosphate utilization
*V*_*c,max*_: Maximum rate of carboxylation

## Supplementary Data

Supplementary Fig.S1. Air temperature and precipitation recorded during the field experiments in Bonn, Cologne, Düsseldorf.

Supplementary Fig.S2. Light response curves of PSII parameters and SPAD for the 23 barley inbred lines.

Supplementary Fig.S3. Developmental changes in heritability.

Supplementary Fig.S4. The growth trajectories for 23 barley inbred lines grown in the field.

Supplementary Fig.S5. Boxplot of the adjusted entry means for 11 morphological traits of twenty-three barley inbred lines in multiple environments across multiple years experiments assigned to four clusters.

Supplementary Fig.S6. Person correlation coefficients calculated between pairs of adjusted entry means based on each developmental phase of 23 barley inbreds for PSII parameters, SPAD and morphological traits collected from multiple environments and years in the field conditions.

Supplementary Fig.S7. The growth trajectory curves for six barley inbred lines grown in the climate chamber condition.

## Acknowledgement

The authors would like to thank Florian Esser and George Alskief (Institute of Quantitative Genetics and Genomics of Plants, Biology Department, Heinrich Heine University, Düsseldorf, Germany) for the field management in Cologne and Düsseldorf. We also thank Onno Muller (Institut für Bio-und Geowissenschaften (IBG), Forschungszentrum Jülich, Jülich, Germany) for organization of the field trial in Bonn. We would like to thank Urte Schlüter (Institute of Plant Biochemistry, Heinrich Heine University, Düsseldorf, Germany) for support with LI-6800. We would like to thank Katharina Luhmer (Universität Bonn, Bonn, Germany) and Monika Bilstein (Landeshauptstadt Düsseldorf, Düsseldorf, Germany) for offering the weather data. The work in BS and SM labs was supported by the DFG through the Cluster of Excellence on Plant Sciences (CEPLAS, EXC2048). WZ thanks the China Scholarship Council for a followship (No.202009655001).

## Author Contribution

BS, SM conceptualized the study, acquired funding and supervised the work; YG, SM, and BS designed the research. YG, MS, LO, WZ performed experiments. YG analyzed the data. YG, SM, and BS interpreted the results. YG, SM, and BS wrote the manuscript.

## Data Availability

Data are publicly available in the manuscript, and in the supplementary information.

